# Structural basis for ubiquitylation by HOIL-1

**DOI:** 10.1101/2022.11.13.516300

**Authors:** Qilong Wu, Marios G. Koliopoulos, Katrin Rittinger, Benjamin Stieglitz

## Abstract

The linear ubiquitin chain assembly complex (LUBAC) synthesises linear Ub chains which constitute a binding and activation platform for components of the TNF signalling pathway. One of the components of LUBAC is the ubiquitin ligase HOIL-1 which has been shown to generate oxyester linkages on several proteins and on linear polysaccharides. Here we describe the crystal structure of a C-terminal tandem domain construct of HOIL-1 comprising the IBR and RING2 domains. The structure adopts an auto-inhibited conformation in which the catalytic cysteine of the RING2 domain is shielded by the adjacent IBR domain. Activation of HOIL-1 is triggered by linear tetra-Ub binding which enables HOIL-1 to mono-ubiquitylate linear Ub chains and polysaccharides. Interestingly, the structure reveals a unique bi-nuclear Zn-cluster which substitutes the second zinc finger of the canonical RING2 fold. We identify the C-terminal histidine of this bi-nuclear Zn-cluster as the catalytic base required for the ubiquitylation activity of HOIL-1. Our study suggests that the unique zinc-coordinating architecture of RING2 provides a binding platform for ubiquitylation targets.

## Introduction

Stimulation of innate and adaptive immune receptors trigger the activation of multiple ubiquitin ligases which orchestrate the initiation of inflammatory gene expression by transcription factor NFκB (Haas *et al*., 2009). Among those E3 enzymes with essential roles in NF-κB activation is the linear ubiquitin chain assembly complex, LUBAC, which attaches linear (or M1-linked) ubiquitin chains to a set of target proteins (Hrdinka and Gyrd-Hansen, 2017). In contrast to canonical ubiquitin chains which are linked by isopetide bonds between lysine residues and the C-terminal glycine of ubiquitin, M1-Ub chains are generated in a head-to-tail fashion by creating a peptide bond between the N-terminal methionine and C-terminus of the two adjoining Ub moieties. LUBAC is a heterotrimeric ligase which consists of the three proteins HOIP, HOIL-1 (also known as RBCK1) and SHARPIN (Gerlach *et al*., 2011; Ikeda *et al*., 2011; Tokunaga *et al*., 2011). Two of the LUBAC components display Ub ligase activity, HOIP and HOIL-1, and both belong to the family of RBR (Ring-between-RING) ligases. Their catalytic activity resides within a specific combination of three domains: a canonical RING1 domain is followed by two zinc coordinating domains termed IBR (In-between-RING) and RING2. The structural and mechanistic features of this RBR catalytic module have been extensively investigated (Walden and Rittinger, 2018; Cotton and Lechtenberg, 2020). RBR ligases recruit Ub charged E2 enzymes via the RING1 domain. The activated ubiquitin is then moved in a transthiolation reaction from the E2 onto a conserved cysteine located in the first zinc-coordinating loop of the RING2 domain to form a thioester intermediate between ubiquitin and the RBR ligase. Substrate binding near the Ub-thioester intermediate allows an aminolysis reaction to occur which attaches Ub to the target protein. Linear Ub chain synthesis by LUBAC is carried out by the ligase component HOIP which binds to an acceptor Ub molecule in the vicinity of the Ub-thioester at the RING2 domain that leads to formation of a peptide bond between the C-terminus of the donor Ub and the N-terminus of the acceptor Ub (Smit *et al*., 2012; Stieglitz *et al*., 2012). The reaction proceeds via a nucleophilic attack from the N-terminal methionine of the acceptor Ub which is assisted by a catalytic base in form of a histidine located near the catalytic cysteine, a mechanism which is widely conserved among RBR ligases (Stieglitz *et al*., 2013). Interestingly, several studies of the LUBAC component HOIL-1 have now demonstrated that its activity deviates from canonical RBR ligases by eliciting a unique functionally. It has been shown that HOIL-1L mono-ubiquitylates several immune signalling proteins such as MyD88 (Myeloid differentiation factor 88), the IL-1 receptor associated kinases IRAK1and IRAK2 by catalyzing formation of an ester bond between the hydroxy group of serine or threonine residues and the C-terminal glycine of Ub. Moreover, ester-linked ubiquitylation was also reported to take place on ubiquitin itself at T12, S20, T22 and T55, which may have important consequences for the catalytic activity of LUBAC (Kelsall *et al*., 2019; Carvajal *et al*., 2021). HOIL-1 mediated ester-linked ubiquitylation of the distal Ub of a linear chain cannot be further extended by HOIP due to a steric hindrance created by the ester bound Ub molecule. The discovery of HOIL-1 mediated mono-ubiquitylation by oxyester linkages on several targets proteins and on ubiquitin chains provides a new layer of complexity for the regulation of immune pathways by the ubiquitin system (Kelsall, 2022). The ester-linked ubiquitylation activity of HOIL-1 does not only play intricate roles in context of LUBAC mediated signalling but is also involved in the seemingly unrelated background of defective glycogen homeostasis. It was recently shown that HOIL-1 can ubiquitylate polysaccharides under *in vitro* conditions by attaching Ub to the hydroxy group of unbranched carbohydrates (Kelsall *et al*., 2022). This intriguing feature provides a rationale for several clinical case studies that have uncovered a genetic link between HOIL-1 deficiencies and glycogen storage diseases (Boisson *et al*., 2012; Krenn *et al*., 2018; Phadke *et al*., 2020; Nitschke *et al*., 2022). Although the discovery of the unique ubiquitylation activity of HOIL-1 represents a major advancement in our knowledge about the diverse functionality of RBR ligases, its structural and mechanistic basis has not been explored. To understand the molecular requirements which enable HOIL-1 to catalyse the attachment of Ub to proteins or carbohydrates by oxyester linkages, we have solved the crystal structure of a C-terminal tandem domain construct comprising the IBR and RING2 domain and identify a unique Zn-cluster as molecular feature required for catalysis.

## Material and Methods

### Protein expression, labelling and purification

All constructs were expressed and purified as described previously (Stieglitz *et al*., 2012). Point mutations were generated using the QuikChange site-directed mutagenesis kit (Stratagene). All plasmids were verified by DNA sequencing. Protein concentrations were determined by UV-VIS Spectrophotometry at 280nm using calculated extinction coefficients. Bovine mono-ubiquitin was purchased from Sigma and further purified by SEC. Ubiquitin R54C was labelled at position R54 by introducing a cysteine by site-directed mutagenesis. Labeling was carried out with Cy5 maleimide mono-reactive dye (Cytiva) according to the manufacturer’s instructions with subsequent purification by SEC.

### Ubiquitylation assays

Assays were performed using 1µM E1, 10 µM UbcH7, 5 µM HOIP 694-1072 and / or 5 <u>µ</u>M HOIL-1, 30 µM Ub, 1µM Ub R54C-Cy5 and 10mM maltoheptaose. Reactions were incubated at 25 °C in 50 mM HEPES pH7.4 with 150 mM NaCl. Samples were taken at 5, 10, 15, 30 and 60 minutes after addition of 5 mM ATP. Reactions were quenched by snap-freezing in liquid nitrogen and analyzed by SDS-PAGE using an iBright Imaging System (Thermo Fisher Scientific) for visualisation of Coomassie Brilliant blue, Lumitein or Cy5.

### Isothermal titration calorimetry

ITC measurements were performed using PEAQ-ITC micro calorimeter (Malvern Panalytical). All samples were dialyzed into buffer containing 50mM HEPES pH 7.5, 150mM NaCl and 1mM TCEP. Titrations were performed at 20 °C with 50 µM of HOIL-1L loaded into the cell and 500 µM linear tetra Ub into the syringe.

### Crystallization, data collection and structure determination

Crystallization trials with HOIL-1 residues 366-510 were set up at 10 mg/ml using an Oryx crystallization robot. Initial hits were optimized by sitting-drop vapor diffusion at 18 °C with a reservoir solution containing 200 mM magnesium chloride hexahydrate, 100mM Tris-HCl pH 7.0, 10% (w/v) PEG 8000. Crystals were flash-frozen in the reservoir solution containing 30% glycerol. Crystals of HOIL-1 (366-510) diffracted to 2.44 Å. A data set was collected on beamline IO2 at the Diamond Light Source (Oxford, UK) and processed using XDS (Kabsch, 2010). The structure was solved by Zn^2+^-SAD phasing. Heavy-atom search, density modification and initial model building was performed using Phenix AutoSol (Adams *et al*., 2010). The model was iteratively improved by manual building in Coot and refined using REFMAC5 and Phenix (Murshudov, Vagin and Dodson, 1997; Emsley and Cowtan, 2004). The stereochemistry of the final models was analyzed with Procheck (Laskowski *et al*., 1993). Data collection and refinement statistics are collated in Table 1. Structural figures were prepared in PyMOL (Schrödinger, L. & DeLano, W., 2020. PyMOL, Available at: http://www.pymol.org/pymol)

## Results

We have previously shown that HOIP is the principal LUBAC component which catalyses the formation of linear Ub chains (Stieglitz *et al*., 2013). In contrast, HOIL-1 displays only poor ligase activity and is dispensable for linear Ub chains synthesis (Smit *et al*., 2012; Stieglitz *et al*., 2012). However, several studies have recently reported that HOIL-1 exhibits a robust ester-linked ubiquitylation activity for linear Ub chains as well as maltoheptaose, which acts as a proxy for unbranched polysaccharides (Kelsall, 2022). Both types of ubiquitylation activities depend on the presence of linear tetra-Ub which binds to HOIL-1L with nanomolar affinity (Fig.1A) and might act as an allosteric activator (Kelsall *et al*., 2022). The modification of maltoheptaose can be readily detected in an *in vitro* reconstitution system which consists of purified components of the ubiquitylation cascade. Ubiquitylation of maltoheptaose is observed after 10 min in the presence of tetra-Ub (Fig.1B, C). To observe ester-linked ubiquitylation of ubiquitin, we have set up an assay which takes advantage of the inhibitory effect of oxyester branched Ub chains on HOIP activity. The isolated RBR module of HOIP displays ubiquitylation activity until the entire pool of mono Ub has been converted into linear poly-Ub chains with molecular weights corresponding to chain lengths of 5 to 12 Ub molecules (Fig.1E). In contrast, the linear chains which are synthesized in the presence of HOIL-1 are much shorter which predominantly consist of chains of 2 to 6 Ub moieties (Fig1F). Even after 60 min the mono Ub pool is not depleted, indicating the presence of HOIL-1 restricts the linear Ub chain synthesis activity of HOIP. This result is in agreement with a recent study which demonstrates that HOIL-1 ubiquitylates tetra-Ub on Thr12 and Thr55 which creates a steric block for further chain elongation by HOIP (Carvajal *et al*., 2021). Thus, the reconstitution of linear chain synthesis in the presence of HOIP-RBR and HOIL-1 allows two competing reactions to occur (Fig.1G): First, HOIP generates linear Ub chains from mono Ub. Once tetra-Ub is formed it can act as an allosteric activator for HOIL-1 which adds a molecule of Ub to linear Ub chains via an oxy-ester linkage, thereby blocking HOIP from further linear chain elongation. Our analysis of HOIL-1 ligase activity is in line with recent studies which demonstrate a distinct functionality for HOIL-1, which deviates from other RBR ligases who catalyse amide-linked ubiquitylation.

**Figure 1.**
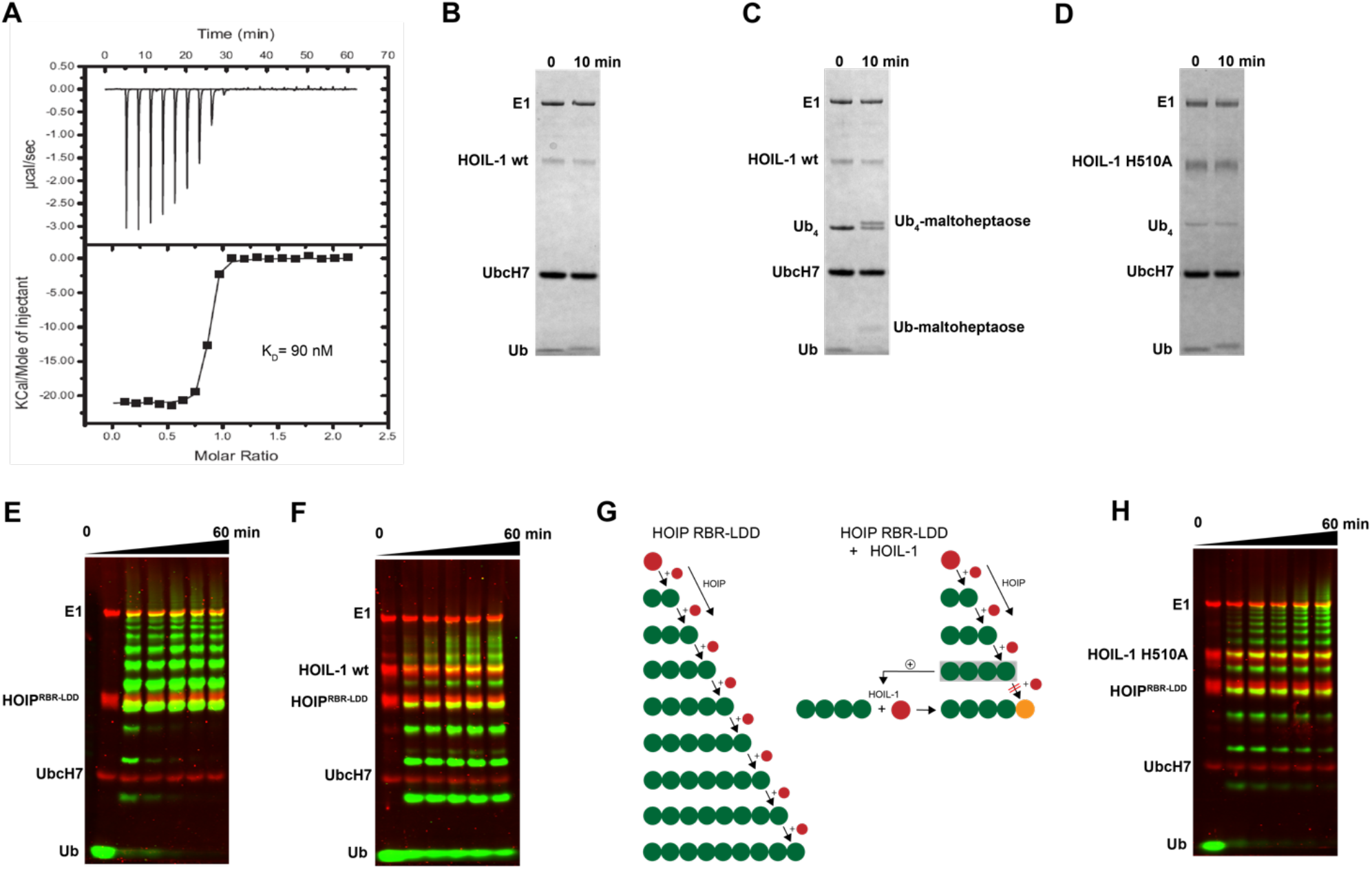
**(A)** Quantitative measurement of HOIL-1 binding to linear Ub_4_ by ITC (K_D_=90 nM). *In vitro* reconstitution of carbohydrate ubiquitylation of HOIL-1 in the absence **(B)** and presence **(C)** of linear tetra Ub. **(D)** The mutation H510A abolishes carbohydrate ubiquitylation of HOIL-1L. **(E)** *In vitro* reconstitution of HOIP-RBR-LDD (residues 697-1072) ligase activity. **(F)** The linear ubiquitin chain synthesis activity of HOIP-RBR-LDD is restricted in the presence of HOIL-1 wt. **(G)** The ubiquitylation activities of HOIP-LDD and HOIL-1 are schematically illustrated with mono-Ub in red, linear Ub in green and ester linked Ub in orange. HOIL-1 is stimulated by linear tetra Ub as indicated by a plus symbol. **(H)** HOIP-LDD mediated linear Ub chain synthesis is re-established in the presence of HOIL-1 H510A.

To understand the molecular determinants for HOIL-1 specific ubiquitylation we analyzed the sequence of HOIL-1 with a focus on the region in immediate vicinity to the catalytic cysteine (Cys460) of the RING2 domain (Fig.2A). A sequence alignment revealed that HOIL-1 together with HOIP, RNF216 and RNF144 deviates in the otherwise conserved pattern of Zn-coordinating residues downstream of the catalytic cysteine. For HOIP and RNF216 this is not surprising since their structures display zinc finger insertions in RING2 which have been shown to be essential for their catalytic activity of M1-linked and K69-linked Ub chain synthesis (Stieglitz *et al*., 2013; Cotton *et al*., 2022). This suggests that HOIL-1 also features a unique Zn-chelating structure within the C-terminal part of RING2 which is potentially important for HOIL-1 catalytic activity. This notion is further supported by a clinical study which reports two patients with amylopectinosis who carry mutations at position 498 or 508 in RING2 of HOIL-1L (Nitschke *et al*., 2022). In addition, unlike most RBR ligases, HOIL-1 does not display a histidine next to the conserved catalytic cysteine which suggests that its catalytic mechanism differs from canonical RBRs. To elucidate the structural features which enable HOIL-1 to exert its unique catalytic activity we solved the crystal structure of fragment 443-510, which comprises the IBR and the RING2 domain (Fig.2B and Table S1). The overall structure shows that both domains are linked by two helices, which are positioned at a 50° angle (Fig.2C). This arrangement brings the IBR and the RING2 domains into close vicinity to each other and allows them to form an interface which buries a total of 1303A^2^ of solvent accessible surface area. The interface is formed by hydrophobic and electrostatic interactions between the N-terminal part of RING2 and the loop regions adjacent to the 4 beta-strands of the IBR domain (Fig.3A). K457 of RING2 is a central contact point for the IBR domain, and is coordinated by F382, E383, V386 and N387. The C-terminal histidine (H510) of RING2 forms a polar interaction with D385 of the IBR domain. These interactions, together with further hydrophobic and polar contributions by D458, K456, Q455, I452, V467 and R464 in RING2 with N387, K403 and L401 of the IBR, create a surface which largely obstructs the catalytic cysteine C460. We therefore assume that the observed structural arrangement represents a catalytic inactive conformation of HOIL-1 in which the catalytic cysteine of RING2 is masked by the IBR domain. As expected from our sequence analysis, the C-terminal part of the RING2 domain of HOIL-1L reveals a unique architecture not observed in other RBR ligases so far: The C-terminal 36 residues form a loop structure which is laced up by a binuclear Zn cluster coordinated by 6 cysteines (Fig.3B). This structure adopts a compact conformation which packs against the second beta sheet of the canonical RING2 fold. HOIL-1 displays a tryptophan (W462) instead of a histidine located two amino acids downstream of the catalytic cysteine (Fig.2A), questioning if HOIL-1 requires a histidine to assist catalysis as observed for other RBR ligases. The loop region of RING2, which wraps around the bi-nuclear Zn-cluster, terminates next to W462. The C-terminal residue of HOIL-1 is a histidine (H510) which forms a pi-staking interaction with W462. In this configuration H510 is ideally positioned to act as a catalytic base. To probe the relevance of H510 for HOIL-1 catalysis, we tested the mutant H510A in our *in vitro* reconstitution assay. Strikingly, this mutant is unable to ubiquitylate maltoheptaose and shows a strongly reduced ability to inhibit HOIP mediated linear Ub chain synthesis, indicating that H510 is required for HOIL-1 catalytic activity (Fig1D,H).

**Figure 2.**
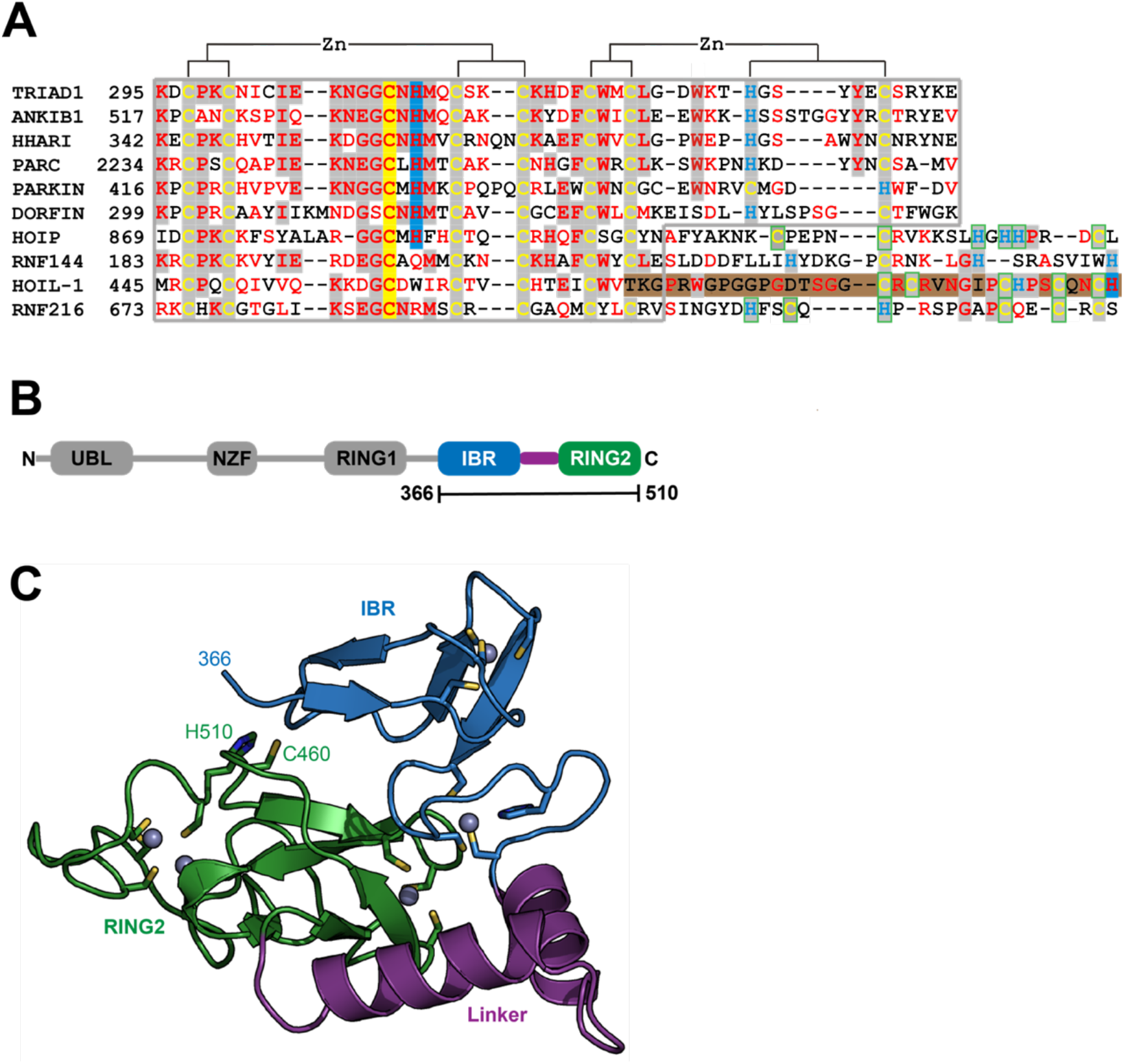
Structural characterisation of HOIL-1 shows a unique deviation of the RING2 domain. **(A)** The RING2 domains of human RBR domains have been aligned according to conserved Zn^2+^ coordinating residues. Catalytic cysteine and histidine residues are depicted on a yellow and blue background, respectively. The C-terminal binuclear Zn-cluster of HOIL-1 is represented on a brown background. **(B)** Domain architecture of HOIL-1 illustrates the boundaries of the crystallized construct. (Domain acronyms: UBL = Ubiquitin Like domain; NZF = Npl4 Zinc Finger; RING1/2 = Really Interesting New Gene domain; IBR = In Between RING domain). **(C)** Structural model of residues 366-510 of HOIL-1 as ribbon models with Zn^2+^ ions as grey spheres. Zn^2+^ coordinating residues, the catalytic cysteine and histidine are shown as a stick model.

**Figure 3.**
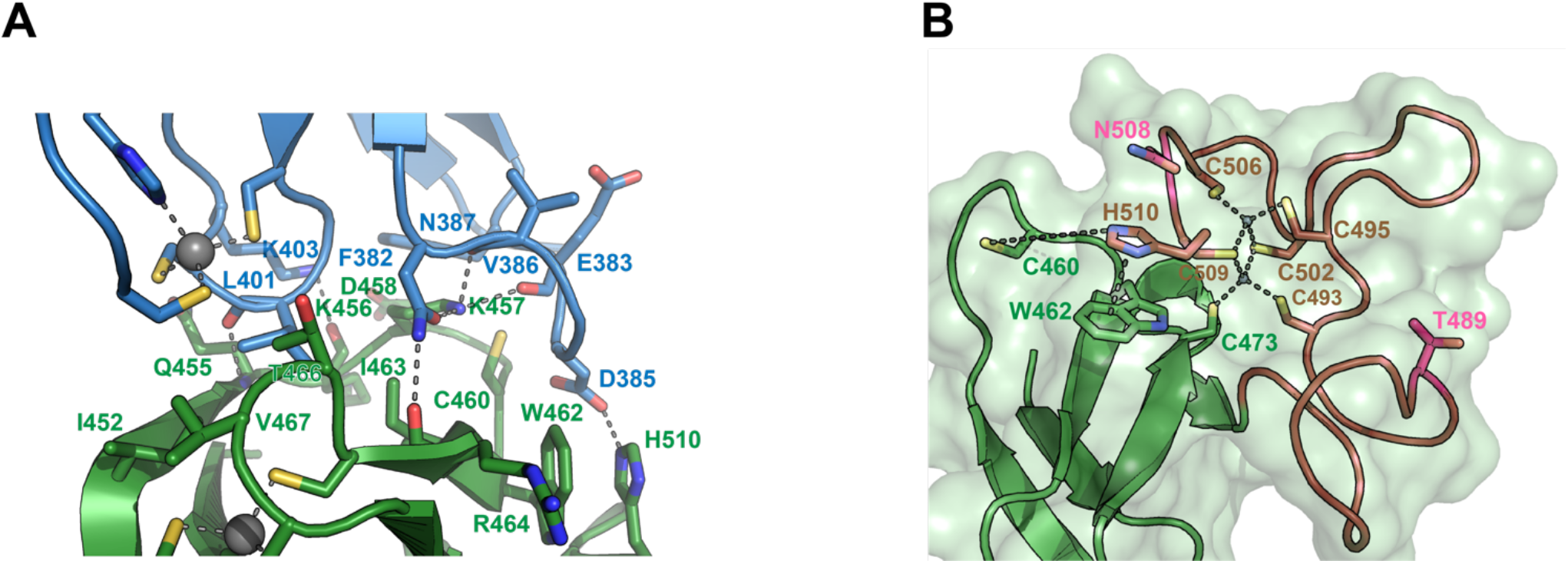
Close-up view of the interface between the IBR domain (blue) and the RING2 domain (green). Residues involved in IBR-RING2 interactions are shown as sticks. The catalytic cysteine (C460) protrudes into a groove of the IBR surface. Polar interactions are shown as dashed lines. The cut-off distance for polar and hydrophobic interactions is 3.5Å and 4Å respectively. **(B)** Close-up view of the C-terminal loop region of RING2 which form the binuclear Zinc-cluster (brown). Zinc coordinating interactions with cysteines are shown as dashed lines. The pi-stacking interaction between H501and W462 is shown as dashed line. The distance between H510 and the catalytic cysteine 460 is 7.6 Å. Positions for pathogenic variants of HOIL-1 (Thr489Profs*9 and Asn508Profs*4) in patients with polyglucosan storage disorders 489 and 508 are shown in pink.

## Discussion

HOIL-1 is a member of the RBR E3 ligase family with the ability to ubiquitylate its targets by linking a hydroxy group of the substrate to the C-terminus of Ub. The recent discovery that the resulting ester bonds can not only be created on proteins but also on carbohydrates, opens new possibilities for the ubiquitin system beyond its definition as a posttranslational modification. Our study sheds light on the structural requirements for the ligase activity of HOIL-1 and uncovers a notable variation of the ubiquitin transfer activity in RBR ligases. Several studies have demonstrated that RBR ligases utilize a histidine as a general base for catalysis, which is located in the first zinc coordinating loop of RING2 next to the catalytic cysteine (Fig.4) (Cotton and Lechtenberg, 2020). HOIL-1 displays a tryptophan in this position but instead provides the catalytic histidine via a unique structural feature not observed in any other RBR ligase. Our structural and biochemical analysis identifies the C-terminal H510 as the catalytic base in HOIL-1. The residue is placed next to the catalytic cysteine through a stretch of 37 residues which fold into a binuclear zinc-cluster. We hypothesize that the extended loop structure of the C-terminal zinc-cluster creates a substrate binding platform for HOIL-1. Specific substrate binding domains have not been identified for any RBR ligase. Based on structural studies on other RBR members such as HOIP, RNF216 and HHARI it becomes increasingly evident that the C-terminal half of RING2 next to the first zinc finger generally determines the substrate specificity in RBR ligases (Stieglitz *et al*., 2013; Cotton *et al*., 2022; Reiter *et al*., 2022; Stieglitz, 2022) (Fig.4). The discovery of the binuclear zinc cluster in RING2 of HOIL-1 supports this hypothesis. The importance of the C-terminal structural architecture for proper HOIL-1 catalysis is also highlighted by the recent identification of mutations in the bi-nuclear Zn cluster in patients with polyglucosan storage disorders (Fig. 3,B) (Nitschke *et al*., 2022). The overall structure of the HOIL-1 tandem domain construct shows that the IBR and the RING2 domains of HOIL-1 are connected by a helical segment. This observation is in-line with the structural models of other RBR ligases, which also display one or two helices to separate both domains from each other. However, in contrast to the structures HOIP, RNF216 and HHARI, which have been captured in their active conformation, the helical linker in HOIL-1 creates a hinge which brings the IBR and the RING2 domain in close contact to each other (Fig.S1) (Lechtenberg *et al*., 2016; Horn-Ghetko *et al*., 2021; Cotton *et al*., 2022). Since this intermolecular interaction partially obstructs the catalytic site of RING2, we speculate that the observed conformation represents a catalytic inactive conformation. In this context it is important to note that inhibition of ligase activity by intramolecular interaction between RING2 and a second domain is a common mechanistic concept in RBR ligases. For example, Parkin is auto-inhibited via interaction of the RING2 domain with its RING0 domain, while HHARI employs the C-terminal Ariadne domain to shield the catalytic cysteine in RING2 (Fig.S2) (Duda *et al*., 2013; Kumar *et al*., 2015). The structure of HOIL-1 might expand this paradigm with the notable variation that no additional domain outside the core RBR module is required to achieve an auto-inhibited conformation. Our biochemical analysis of HOIL-1 enzymatic function shows that the ligase requires the presence of linear tetra Ub for its activity. A previous study has shown that HOIL-1 binds linear and K63 linked Ub chains for allosteric activation and suggested that binding of allosteric Ub chains involve the IBR domain, similar to the allosteric Ub binding site observed in the structure of activated HOIP (Fig.S3) (Kelsall *et al*., 2022). It is tempting to speculate that binding of allosteric poly Ub at this position could dislodge the RING2 domain from the backside of the IBR domain in order to adopt a catalytic competent conformation.

**Figure 4.**
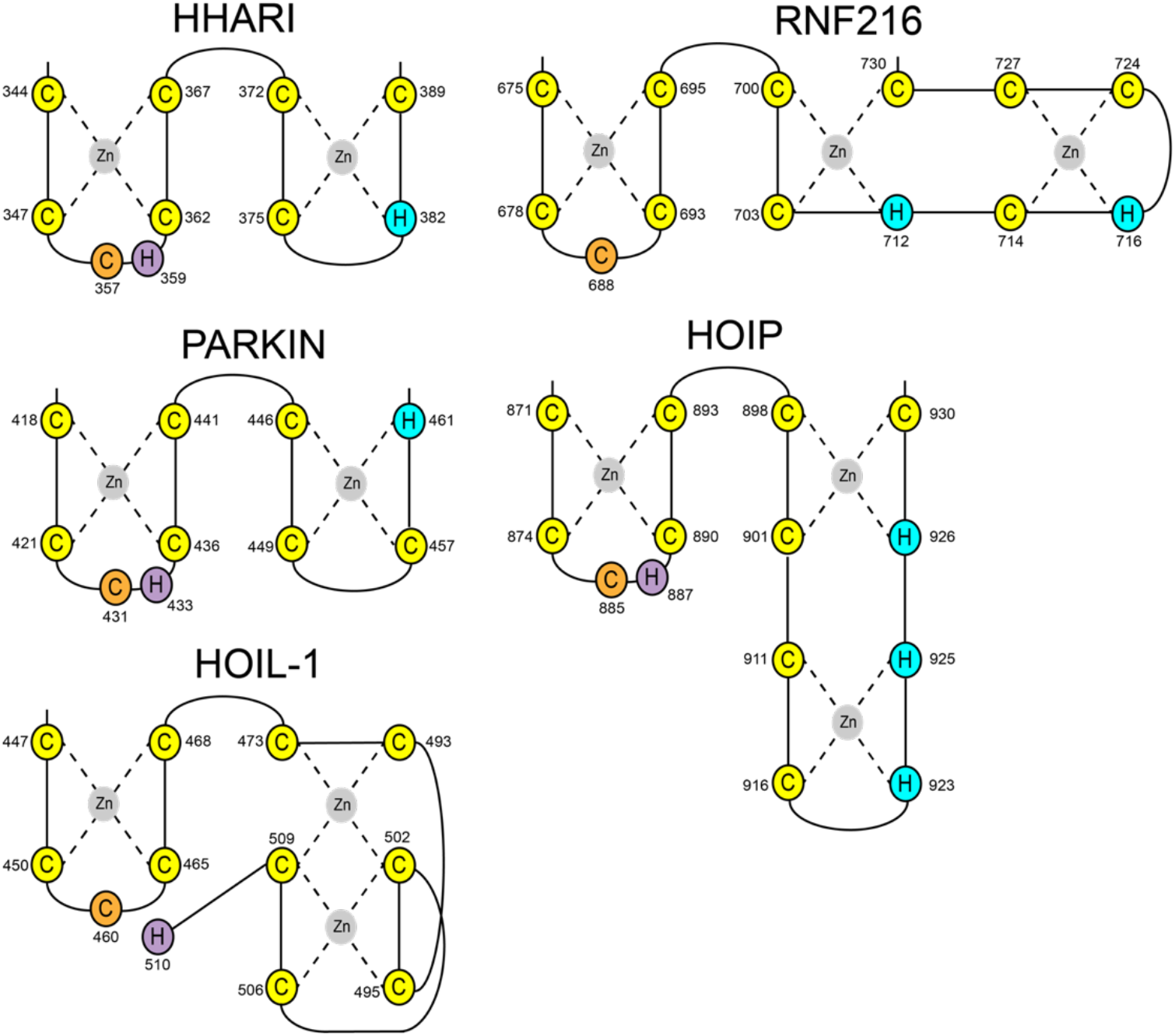
Schematic presentation of RING2 topologies of the RBR ligases HHARI, Parkin, HOIP, RNF216 and HOIL-1. Zinc coordinating cysteine and histidine residues are shown in yellow and blue, respectively. Catalytic cysteines are shown in orange. Catalytic histidine residues have been identified for HHARI, Parkin, HOIP and HOIL-1 (magenta).

In summary, we have uncovered that HOIL-1 adopts a unique structure in form of a binuclear zinc-cluster which is required for its catalytic function. To further dissect the mechanism of HOIL-1 catalysis it will be essential to obtain high-resolution structures of HOIL-1 in its active conformation bound to its substrates. Future studies need to provide an understanding how HOIL-1 is able to ubiquitylate carbohydrates in molecular detail.

## Author Contributions

KR and BS conceived the study and wrote the manuscript. QW, MGK and BS performed and analyzed experiments.

## Acknowledgements

We thank E. Christodoulou for technical assistance, L. Haire for help with crystallization P. Walker for data collection and the Diamond Light Source for synchrotron access.

## Supplementary Material

### 1 Supplementary Figures and Tables

#### 1.1 Supplementary Figures

**Supplementary Figure 1.**
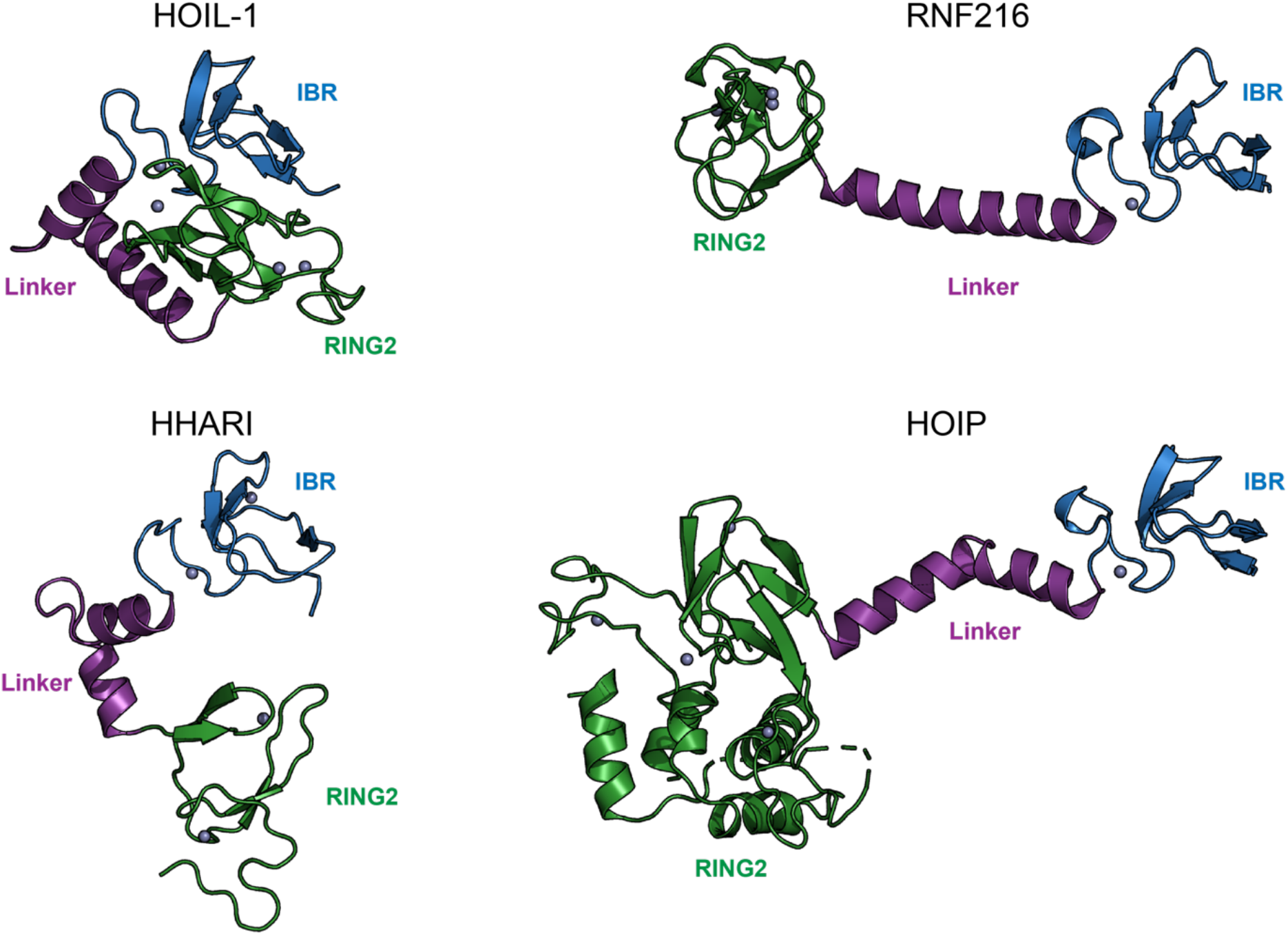
Comparison of IBR-RING2 orientations in RBR ligases. The structure of HOIL-1 is compared with the IBR-RING2 conformations of the activated ligases RNF216 (PDB: 7M4M), HHARI (PDB: 7B5L) and HOIP (PDB: 5EDV). Structures are shown as ribbon models with Zn^2+^ ions as grey spheres.

**Supplementary Figure 2.**
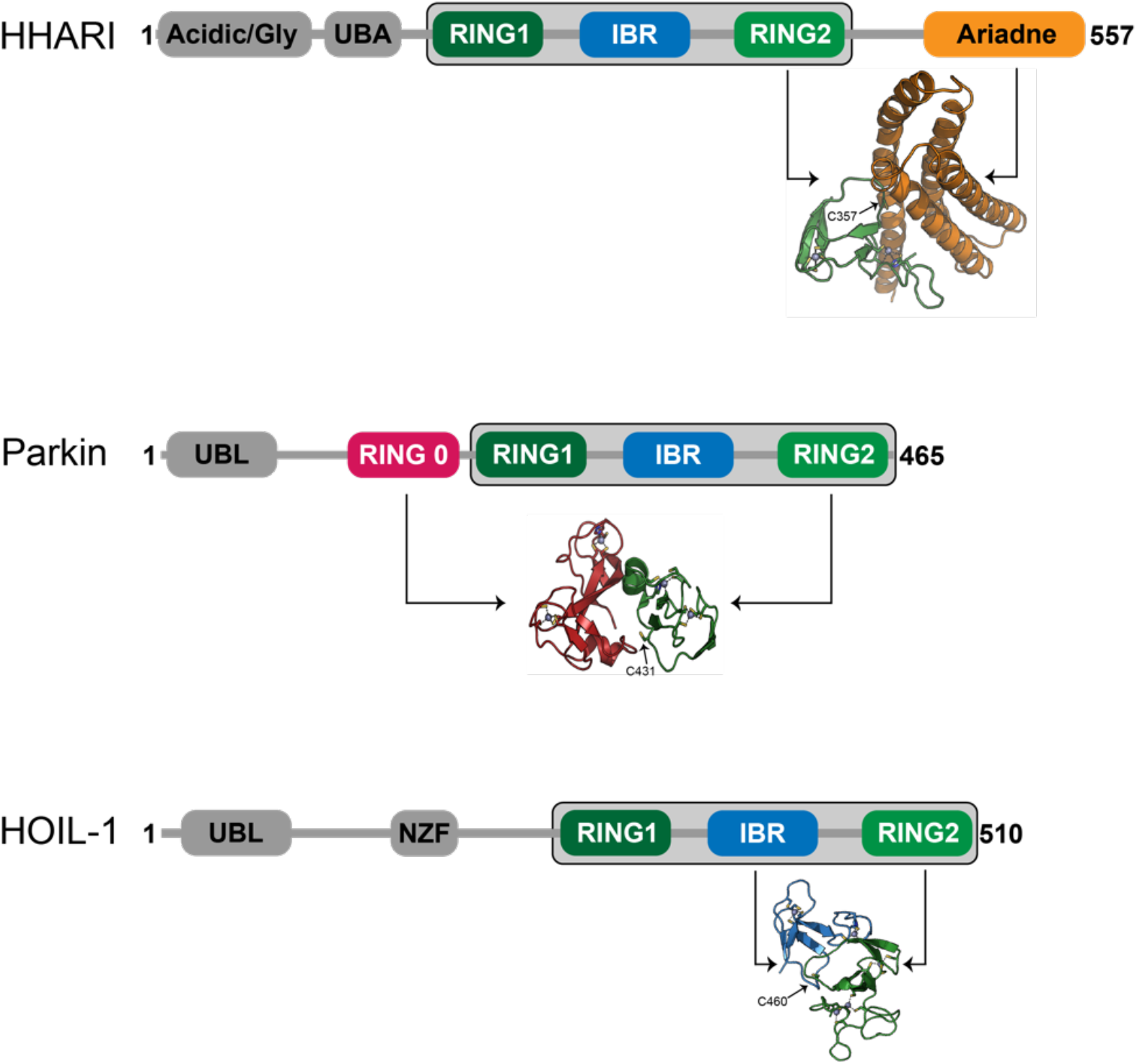
Domain architecture and structural comparison of intramolecular RING2 interactions of HHARI, Parkin and HOIL-1. The RBR core module is indicated by a grey box. (Domain acronyms: UBL = Ubiquitin Like domain; NZF = Npl4 Zinc Finger; UBA = Ubiquitin Associated domain; Acidic/Gly = Acidic and Glycine rich domain; Ariadne = Ariadne domain; RING0/1/2 = Really Interesting New Gene domain; IBR = In Between RING domain) The intramolecular complexes between RING2 and corresponding domains are shown as a ribbon model. The Zn^2+^ coordinating residues and catalytic cysteines of HOIL-1, HHARI (PDB: 4KBL) and Parkin (PDB: 5C1Z) are indicated. Zn^2+^ ions are shown as grey spheres.

**Supplementary Figure 3.**
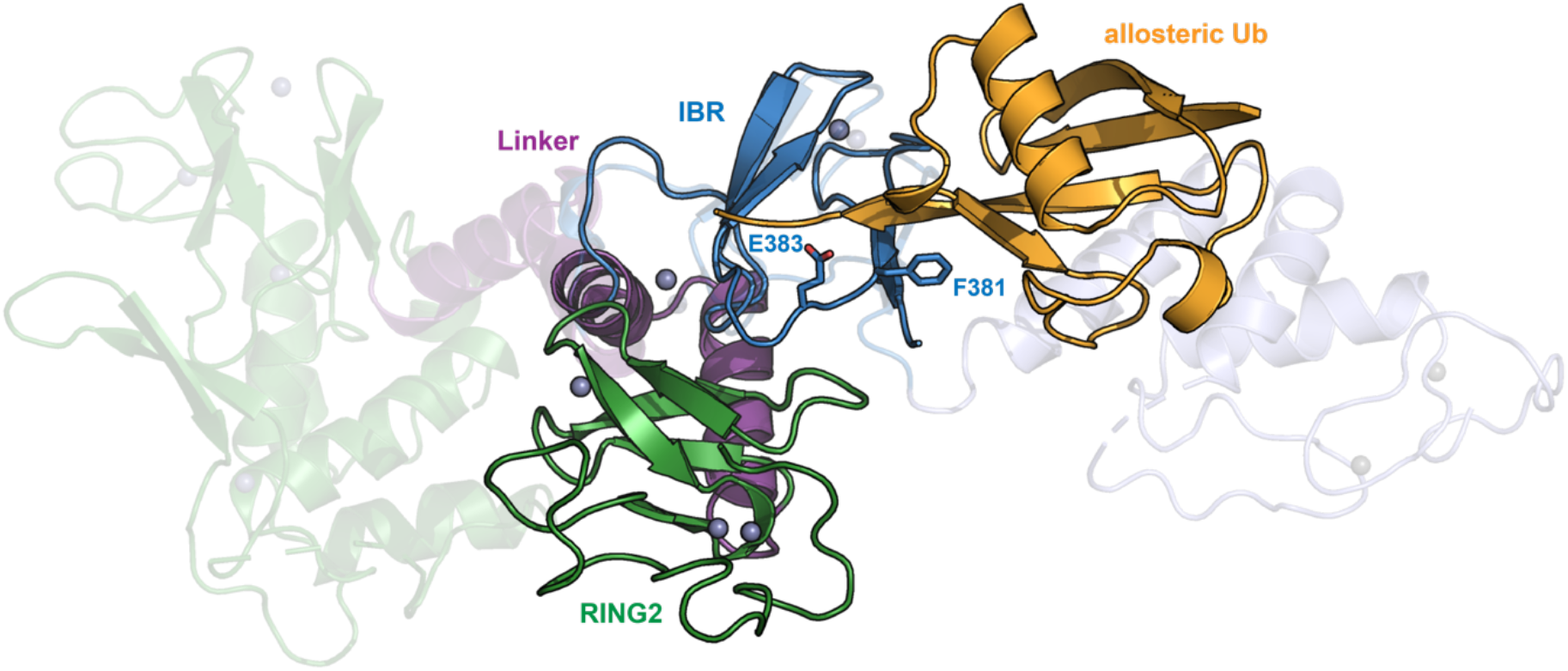
Structural overlay of HOIL-1 IBR-RING2 with the active conformation of HOIP (PDB: 5EDV; translucent ribbon model) in complex with ubiquitin (orange). The structures are superimposed over the entire length of the IBR domains. Residues F381 and E386 are essential for HOIL activation (indicated as ball-and stick model).

#### 1.2 Supplementary Table

**Table S1.** Data collection and refinement statistics.

| Data collection |  |
| --- | --- |
| Space group | P 1 2<sub>1</sub> 1 |
| Cell dimensions |  |
| a, b, c (Å) | 53.91, 59.25, 57.21 |
| $\alpha$, $\beta$, $\gamma$ (°) | 90.00, 90.31, 90.00 |
| Wavelength (Å) | 1.2829 |
| Resolution range (Å) | 41.16 - (2.32 - 2.24) |
| Total reflections | 33358 (3291) |
| Unique reflections | 16782 (1651) |
| Completeness (%) | 95.79 (95.81) |
| $I / \sigma I$ | 10.14 (3.75) |
| $R_{\text{merge}}$ | 0.08046 (0.2176) |
| $CC_{1/2}$ | 0.989 (0.914) |
| Refinement |  |
| Resolution range (Å) | 41.16 - 2.24 |
| Reflections | 16771 |
| $R_{\text{work}}^a / R_{\text{free}}^b$ | 24.73/29.45 |
| Number of atoms | 2342 |
| Protein | 2254 |
| Zn<sup>2+</sup> | 10 |
| Water | 78 |
| $B$-factors average (Å<sup>2</sup>) | 40.41 |
| Protein (Å<sup>2</sup>) | 40.77 |
| Zn<sup>2+</sup> (Å<sup>2</sup>) | 26.16 |
| Water (Å<sup>2</sup>) | 31.57 |
| R.m.s. deviations |  |
| Bond lengths (Å) | 0.008 |
| Bond angles (°) | 1.00 |
| Ramachandran |  |
| favoured, allowed, outliers (%) | 93.71, 4.90, 1.40 |

Statistics for the highest-resolution shell are shown in parentheses.
<sup>a</sup> $R_{\text{work}} = \sum_{\text{hkl}} ||F_{\text{obs}}(\text{hkl})| - |F_{\text{calc}}(\text{hkl})|| / \sum_{\text{hkl}} |F_{\text{obs}}(\text{hkl})|$ .
<sup>b</sup> $R_{\text{free}}$ = the cross-validation $R$ factor for 5% of reflections against which the model was not refined.

